# Filamentous calcareous alga provides a substrate for coral-competitive macroalgae in the degraded lagoon of Dongsha Atoll, Taiwan

**DOI:** 10.1101/363796

**Authors:** Carolin Nieder, Chaolun Allen Chen, Shao-Lun Liu

**Author notes:** Corresponding author; (SLL).

## Abstract

**Background:** The chemically-rich seaweed *Galaxaura* is not only highly competitive with corals, but also provides substrate for other macroalgae. Its ecology and associated epiphytes remain largely unexplored. To fill this knowledge gap, we herein undertook an ecological assessment to explore the spatial variation, temporal dynamics, and epiphytic macroalgae of *G. divaricata* on patch reefs in the lagoon of Dongsha Atoll, a shallow coral reef ecosystem in the northern South China Sea, repeatedly impacted by mass coral bleaching events.

**Methods:** Twelve spatially independent patch reefs in the Dongsha lagoon were first surveyed to assess the benthic composition in April 2016, and then revisited to determine *G. divaricata* percent cover in September 2017, with one additional *Galaxaura*-dominated reef (site 9). Four surveys over a period of 17 months were carried out on a degraded patch reef (site 7) to assess the temporal variation in *G. divaricata* cover. Epiphytic macroalgae associated with *G. divaricata* were quantified and identified through the aid of DNA barcoding.

**Results:** Patch reefs in the Dongsha lagoon were degraded, exhibiting relatively low live coral cover (5-43%), but high proportions of macroalgae (13-58%) and other substrates (rubble and dead corals; 23-69%). The distribution of *G. divaricata* was heterogeneous across the lagoon, with highest abundance (16-41%) in the southeast area. Temporal surveys from site 7 and photo-evidence from site 9 suggested that an overgrowth by *G. divaricata* was still present to a similar extend after 17 months and 3.5 years. Yet, *G. divaricata* provides a suitable substrate some allelopathic macroalgae (e.g., *Lobophora* sp.).

**Conclusions:** Our study demonstrates that an allelopathic seaweed, such as *G. divaricata*, can overgrow degraded coral reefs for extended periods of time. By providing habitat for harmful macroalgae, a prolonged *Galaxaura* overgrowth could strengthen negative feedback loops on degraded coral reefs, further decreasing their recovery potential.

## Introduction

Coral-macroalgae competition is a naturally ecological process on coral reefs [1]. However, anthropogenic disturbances, e.g., climate change, overfishing, and pollution, have intensified space competition of macroalgae against corals and in turn led to a phase shift from a coral-dominated to a macroalgae-dominated ecosystem [2]. The recovery of live corals on degraded reefs is strongly influenced by the types of dominant macroalgae, i.e., allelopathic versus non-allelopathic types [3,4].

Allelopathic macroalgae produce lipid-soluble secondary metabolites, e.g., loliolide derivatives or terpenes, that are poisonous to corals (known as allelochemicals). Such allelochemicals are capable of bleaching and killing coral tissue [5], decreasing the photosynthetic efficiency of zooxanthellae [6], and altering the coral microbiome, ultimately decreasing coral health [7,8]. Allelopathic macroalgae are considered most detrimental for the resilience of coral reefs [12], as these types may perpetuate their dominance by deterring coral larval settlement, and inhibiting the growth and survival of juvenile recruits, key processes of coral reef recovery [9–11].

The red upright calcifying seaweed *Galaxaura* is known to be highly allelopathic against corals. Life history of the genus *Galaxaura* can be grouped into two morphotypes, a smooth and a filamentous type. The latter is characterized by hairy branches that are covered with fine assimilatory filaments [12]. Extracts of the lipid-soluble secondary metabolites of *G. filamentosa* were shown to cause bleaching and death of coral tissue [13,14], and deterred coral larvae from settling [15]. It has thus been suggested that high abundance of *Galaxaura* on degraded reefs can inhibit the recovery of live coral cover [4,15,16].

The filamentous morphotype of *G. divaricata*, is widely distributed in subtropical and tropical reef areas in the Pacific Ocean [17]. Filamentous *G. divaricata* is also common on coral reefs in the shallow lagoon of Dongsha Atoll [18]. Dongsha Atoll is the only large (> 500 km^2^) coral reef atoll in the northern South China and represents a highly valuable hot-spot for marine biodiversity in this region [19]. A catastrophic mass bleaching in 1998 and reoccurring bleaching events thereafter have, however, caused severe mass mortalities of corals in the Dongsha lagoon, followed by a marked increase of macroalgae [20,21]. To date, little is known about the current state of recovery and dominant macroalgae in Dongsha lagoon patch reefs. The proliferation of *G. divaricata* on degraded reefs in the lagoon of Dongsha Atoll was first uncovered during a systematic macroalgae sampling expedition in February 2014 [18]. Of interest to us was our observation that *G. divaricata* was highly populated by macroalgae. Habitat formation is known from other macroalgae (e.g., crustose *Lobophora* or canopy-forming *Sargassum* and *Turbinaria*) that provide substrate for epiphytic algae [22–24]. The dense epiphytic community associated with *G. divaricata* might indicate a previously unappreciated role of *Galaxaura* as a habitat forming seaweed.

The goals of this study were to 1) assess the benthic composition of lagoon patch reefs, 2) document the spatial distribution of *G. divaricata* on patch reefs in the lagoon, 3) monitor the temporal dynamics of *G. divaricata* percent cover over time, and 4) quantify and identify the epiphytic macroalgae associated with *G. divaricata*. The provision of new habitat for other macroalgae by *G. divaricata* could have several ecological implications, worth exploring. For instance, the epiphytic community on *G. divaricata* may enhance macroalgae biodiversity on the reef, or provide trophic support for herbivores, while a facilitation of allelopathic algal types would decrease the resilience of coral reefs.

## Materials and methods

### Ethics statement

The ecological assessments and sample collections in this study were conducted with permissions of the Dongsha Atoll National Park.

### Site description

This study was conducted from April in 2016 to September 2017 in the lagoon of Dongsha Atoll (also known as Pratas Island; 20°40’43” N, 116°42’54” E), which is an isolated coral reef atoll in the northern South China Sea. The atoll covers an area of approximately 500 km^2^ and is situated 450 km southwest from the coast of Taiwan and 350 km southeast from Hong Kong (Fig 1A). The climate is seasonal and varies between a northeast monsoon winter (October-April) and southwest monsoon summer (May-September) [25]. Field work during the northeast winter monsoon is often restricted due to local weather conditions. The ring-shaped reef flat encircles a large lagoon with seagrass beds and hundreds of coral patch reefs [26]. Channels, at the north and south of the small islet (1.74km^2^), interrupt the reef flat, allowing for water exchange between the lagoon and the open ocean. The semi-closed lagoon is about 20 km wide with a maximum depth of 16 m near the center [20]. The Lagoon patch reefs are structured into reef tops (1-5 m depth) and reef slopes (5-12 m depth), and provide important habitat and sheltered nursery grounds for numerous marine organisms, such as green sea turtles and coral reef fish, including rays and sharks [26]. For background information the lagoon water temperature was measured at each survey site, every 30 min from March 2016 to September 2017 using HOBO Pendant^®^ Temperature/Light 8K Data Loggers (UA-002-08, Onset Computer Corporation, USA). Water temperatures were highest during the summer monsoon, averaging 30.1°C, and lowest during the winter monsoon, averaging 24.8°C. Maximum temperatures from July to August reached 34°C on reef tops and 32.7°C on reef slopes.

**Fig 1.**
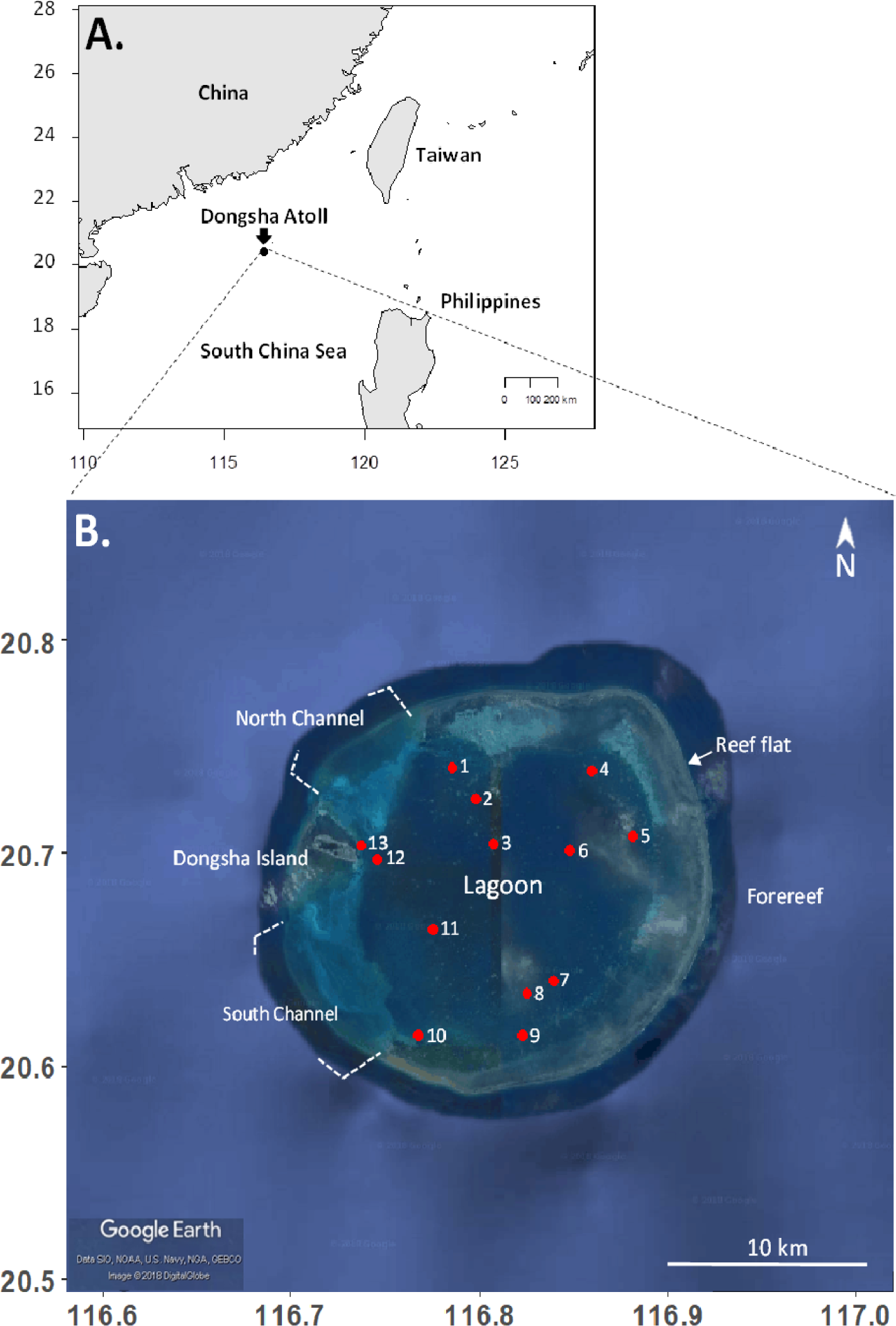
Study site. A) Geographical location of Dongsha Atoll in the northern South China Sea. B) Lagoon patch reef sites surveyed in this study.

### Spatial variation in benthic composition and *G. divaricata* cover of lagoon patch reefs

To assess the benthic composition of patch reefs in the lagoon of Dongsha Atoll, 12 spatially independent reefs were initially surveyed with SCUBA in April 2016 (Fig 1B and S1 Table). A 45-m transect was laid out across each reef area: reef top (1-5 m depth) and reef slope (5-12 m depth). The two transects were 10-20 m apart from each other. The percent cover of live corals, total macroalgae (MA; all upright growing (including *G. divaricata*) and crustose non-coralline seaweeds, and low growing, filamentous turf algae [27]), crustose coralline algae (CCA), and other substrates was estimated using a 35 cm x 50 cm PVC sapling frame [28]. Other substrates mainly constituted dead coral skeleton, rubble, and rocks covered with sediments. Estimates were done *in-situ* at every meter mark, with a total of 45 sampling frames analyzed per transect. The 12 sites were revisited in September 2017 to estimate the percent cover of *G. divaricata* and live corals only, using the same survey method described above. An additional patch reef (site 9) was included, as this site was historically shown to be dominated by *G. divaricata* based on photo evidence, resulting in a total of 13 survey sites (Fig 1B and S1 Table). The diameter of haphazardly selected *G. divaricata* thalli were measured *in situ* at each site and classified as small (1-5 cm diameter), medium (>5-15 cm diam.), and large (>15-30 cm diam.).

### Temporal variation in *G. divaricata* cover

To assess variations in the *G. divaricata* cover over time, we selected the slope area of a degraded patch reef (site 7) that was considerably overgrown by *G. divaricata* (14-18%) and had relatively low coral cover (13-19%). Percent cover of *G. divaricata*, and live corals were estimated in April 2016, the last month of the winter monsoon season, and three times in the summer monsoon season in July, and September 2016, and in September 2017, spanning a period of 17 months. At each time 45 photographs were taken in 1 m intervals along a 45 m fixed transect with an Olympus Stylus-TOUGH TG4 digital camera (25-100 lens, 35mm equivalent) mounted onto a PVC-quadrat (height = 0.64 cm) above a 35 cm x 50 cm sampling frame. Cover estimates were obtained from photographs using ImageJ software, and a superimposed 10 x 10 reference grid, where 1 square represented 1 % of the total grid area. *G. divaricata* cover estimates were arbitrarily ranked into four different categories: very low (0-1.5%), low (>1.5 – 5%), high (>5-20%), and very high (>20%).

### Epiphytic macroalgae associated with *G. divaricata*

This study was carried out in September 2017. Thirty thalli of *G. divaricata* were collected from a degraded reef (site 7) with relatively high percent cover of *G. divaricata* (14-18%). *G. divaricata* thalli were haphazardly collected along a 45-m transect at 5 m depth. Epiphytic macroalgae were removed and identified to the closest identifiable taxonomic unit, using either the Dongsha seaweed guide book [18] or DNA barcoding. The presence and absence of each taxonomic unit was recorded, and the occurrence frequency (*f*) was calculated as follow: *f* = *c (taxonomic unit*_*i*_)/*n*, where *c (taxonomic unit*_*i*_) stands for the count number of thalli that have the epiphyte taxonomic unit *i*, and *n* equals 30, the total number of thalli analyzed. For DNA barcoding, macroalgae samples were preserved in silica gel after collection, and the total genomic DNA of samples was extracted with Quick-DNA™ Plant/Seed Miniprep Kit (Zymo Research Co., USA). Primers for the plastid gene specific amplifications were used as follows: *rbc*L *F7/R753* for red algae [29], *rbc*L *F68/R708* for brown algae [30], and *tufA F210/R1062* for green algae [31]. The newly generated sequences were deposited in GenBank and searched using BLASTn against the GenBank database (S2 and S3 Tables). Sequence similarities of >98% were considered for species identification.

### Statistical analysis

Spatial variations in the percent cover of major benthic categories (corals, total macroalgae, crustose coralline algae, and other substrates) and *G. divaricata* were compared between two reef areas (top and slope) among sites using a two-way ANOVA, with area and site as fixed factors. Similarly, a two-way ANOVA was applied to evaluate the temporal variations in the cover of two major benthic categories (live corals and *G. divaricata*) among four time points, with benthic category and time as fixed factors. A significant difference was considered for p-values lower than 0.05. Maps and statistical graphs were done using R software.

## Results

### Benthic composition

Our spatial survey showed that both live coral cover and total macroalgae cover significantly varied between reef top (1-5 m) and reef slope (5-10 m) and among sites (area × site: F_11, 1056_ = 17.601-26.27, *P* < 0.05; Figs 2A and 2B, and S4 Table). Percent cover for corals, macroalgae, and CCA were generally higher on the reef top, while other substrates were slightly higher on the slope (area: F_1, 1056_ = 6.617-62.725, *P* < 0.05; Figs 2A-2C and S4 Table). We found that the macroalgae cover generally exceeded live coral cover on patch reefs in the Dongsha lagoon (Figs 2A and 2B). Using an arbitrary cutoff of 25%, we observed a higher coral cover in the west of the lagoon, in between the North and South channel, where water exchange is more efficient, i.e., sites1-3, 8, 11, and 13 (Fig 2A). In contrast, no clear spatial pattern of the macroalgae cover was observed. Using a 50% cutoff, it, however, appeared that the shallow and calm area in the southeast lagoon showed a higher macroalgae cover than other areas (i.e., sites 7 and 10; Fig 2B). Compared with live corals and total macroalgae, the CCA cover was relatively low (range: 1-3%; Fig 2C), while the average “other substrates” cover (mainly dead coral skeletons, rubble, and rocks) was extremely high (range: 23-69%; Fig 2D).

**Fig 2.**
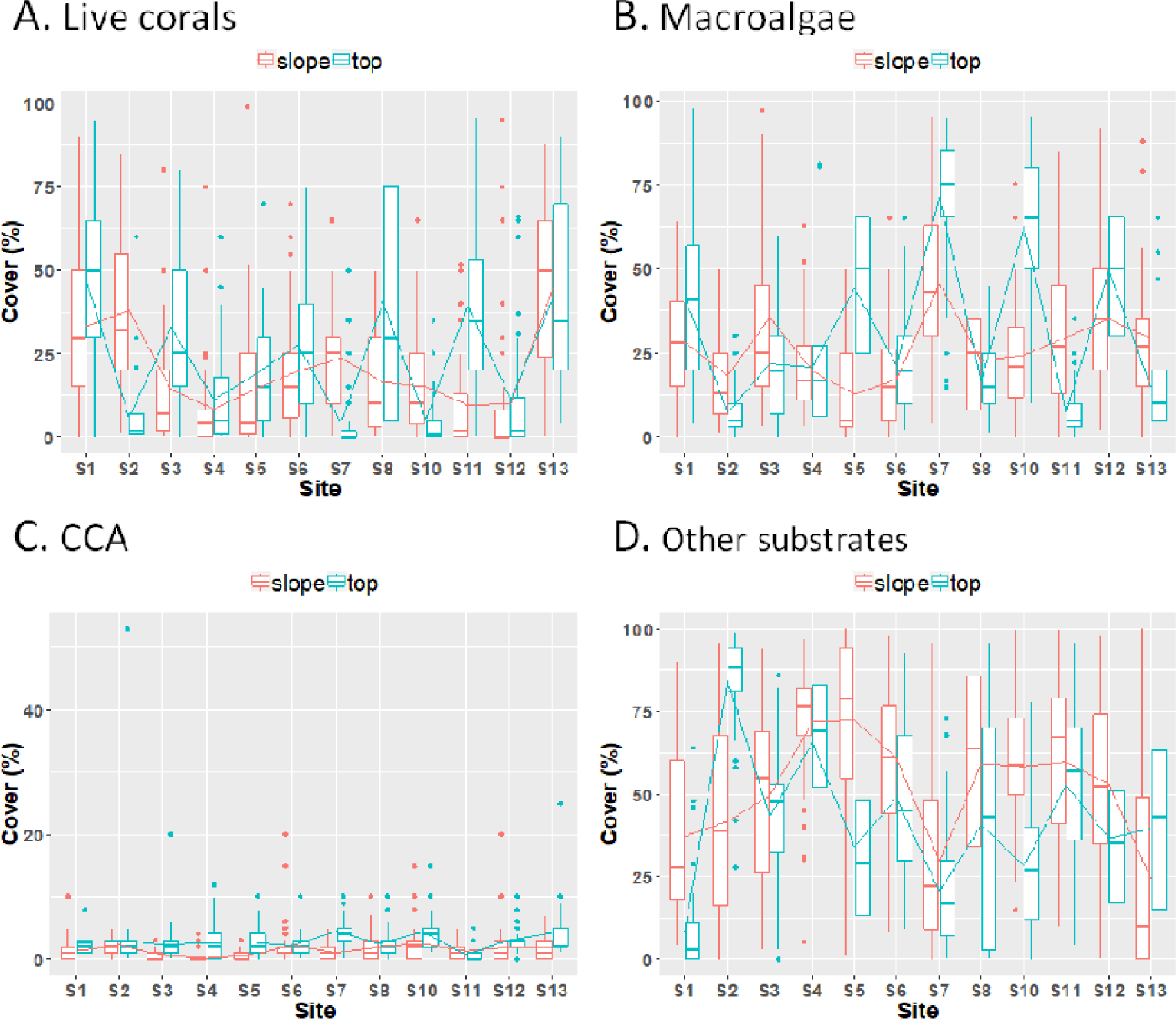
Spatial variation in benthic categories in the Dongsha lagoon. Variation in the cover of (A) live corals, (B) total macroalgae (upright and crustose non-coralline seaweeds, and turf), (C) crustose coralline algae (CCA), and (D) other substrates (dead coral skeletons, rubble, and rocks) between two reef areas (top and slope) among 12 sites.

### Spatial variations in *G. divaricata* cover

The percent cover of *Galaxaura divaricata* was significantly different between reef tops and reef slopes, showing a higher percent cover on the slope (area: F_1, 1144_ = 6.574, *P* < 0.05; S5 Table). There was a significant statistical interaction between area (slope and top) and site (e.g., top > slope at site 5 and slope > top at site 7; area × site: F_12, 1144_ = 7.460, *P* < 0.05; Fig 3 and S5 Table). *G. divaricata* cover was significantly different among the 13 sites (site: F_12, 1144_ = 179.278, *P* < 0.05; Fig 3 and S5 Table), showing highest cover in the southeast area of the lagoon, i.e., site 9 (41%) and the slope of site 7 (16%) (Fig 3). Patch reefs in the northeast lagoon exhibited moderate, low, and very low cover of *G. divaricata* (range: 0.21-5.7%) (Fig 3 and S6 Table). Survey sites in the south, center, west, and north of the lagoon were characterized by very low cover of *G. divaricata* (range: 0-1.4%; Fig 3 and S6 Table). During our survey, we observed that the thallus shape and size of *G. divaricata* varied across sites (S1 Fig). Small ball-shaped or slender thalli were dominant on patch reefs in the northeast lagoon, while medium ball-shaped and large, carpet-like thalli were exclusively present in the southeast lagoon. To further rule out the possibility of cryptic species, our DNA barcoding analyses confirmed that all samples across sites were 100% identical in their *rbc*L sequences, indicative of conspecificity (S3 Table).

**Fig 3.**
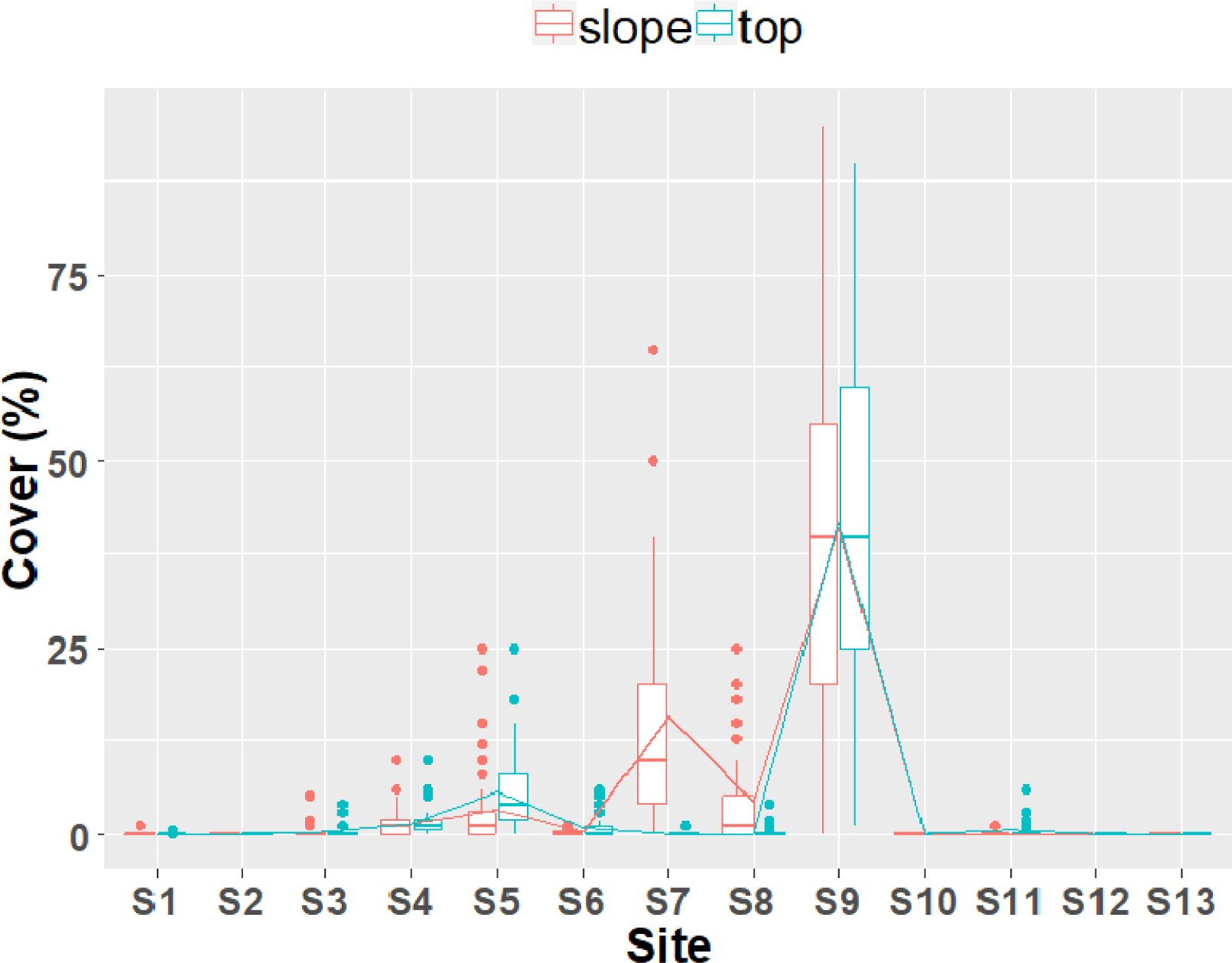
Spatial variation in *G. divaricata* in the Dongsha lagoon. Variation in the cover of *G. divaricata* between two reef areas (top and slope) among 13 sites.

### Temporal dynamics of *G. divaricata* cover

Our temporal survey at a *Galaxaura*-dominated reef (the slope area of site 7) revealed that the percent cover of *G. divaricata* did not vary significantly among the surveys conducted at four time points (April 2016, July 2016, September 2016, and September 2017) over a period of 17 months (time: F_3, 352_ = 0.632, *P* = 0.595; Fig 4 and S7 Table). The percent cover between live corals and *G. divaricata* did not differ significantly (benthic category: F_1, 352_ = 0.086, *P* = 0.770; Fig 4 and S7 Table). Overall, there was no significant statistical interaction between the percent cover of *G. divaricata* and live corals among time points (benthic category × time: F_3, 352_ = 0.363, *P* = 0.780; Fig 4 and S7 Table). Across four time points the mean *G. divaricata* cover remained relatively high (16.45 ± 1.17%), while mean coral cover was low (15.91 ± 0.6%). In addition, we provide photo-evidence from an additional patch reef (site 9, 3-5 m) overgrown by *G. divaricata*. Photographs of the site were taken in February 2014 and in September 2017, showing that the same *G. divaricata* overgrowth was present after 3.5 years (Figs 5A and 5B). *G. divaricata* frequently grew on live corals, where the holdfast penetrated the calcium-carbonate structure, creating a strong attachment to the corals (Fig 5C). In several cases we observed a fluorescent pink discoloration and bleaching of the coral tissue at the contact zone with *G. divaricata*, strongly indicative of allelopathic inhibition by *G. divaricata* (Fig 5D).

**Fig 4.**
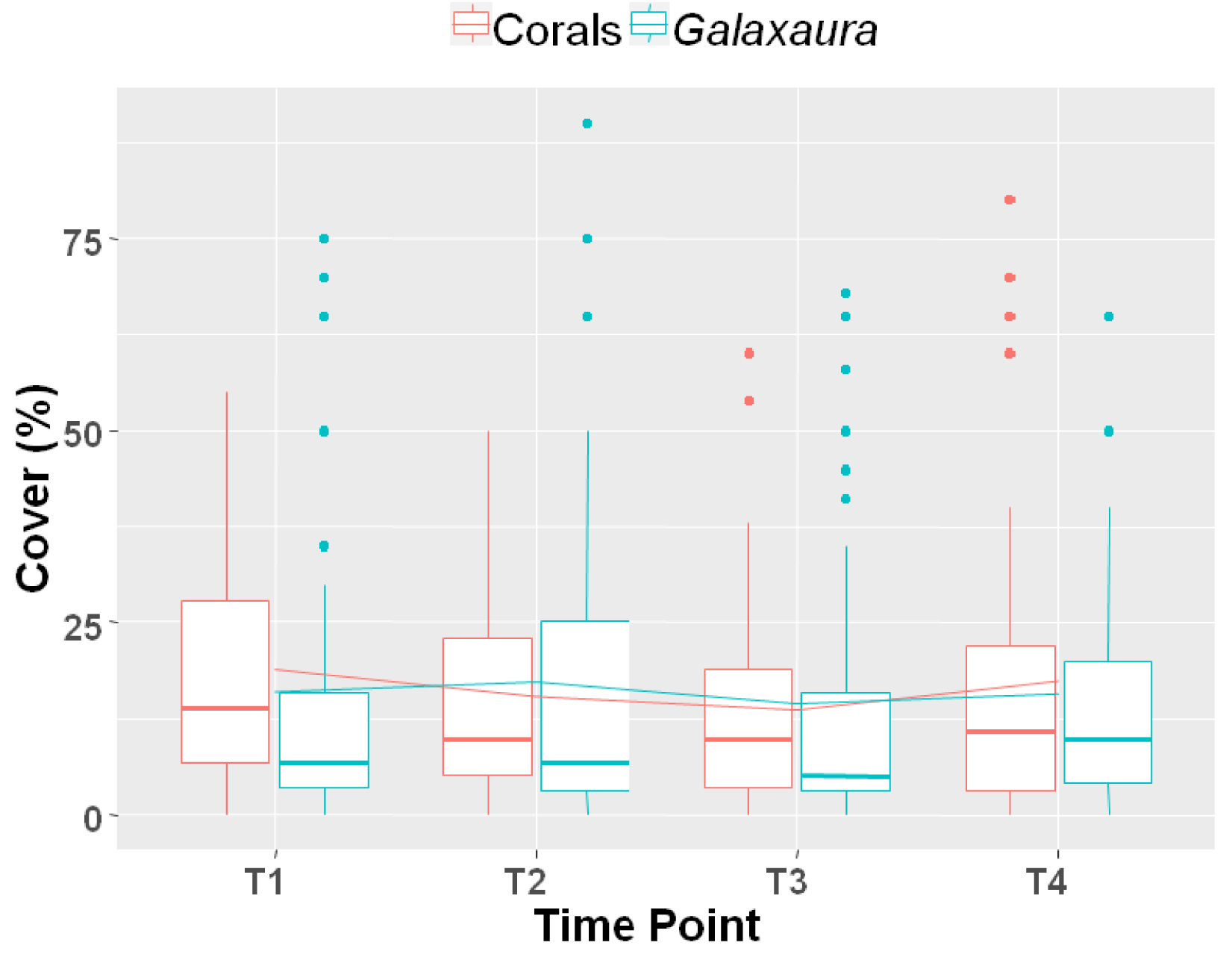
Temporal variation in *G. divaricata* on a degraded patch reef in the southeast Dongsha lagoon. Variation in the percent cover of two major benthic categories (live corals and *G. divaricata*) among four time points (T1: April 2016, T2: July 2016, T3: September 2016, and T4: September 2017) over a period of 17 months (about 5 m depth at site 7).

**Fig 5.**
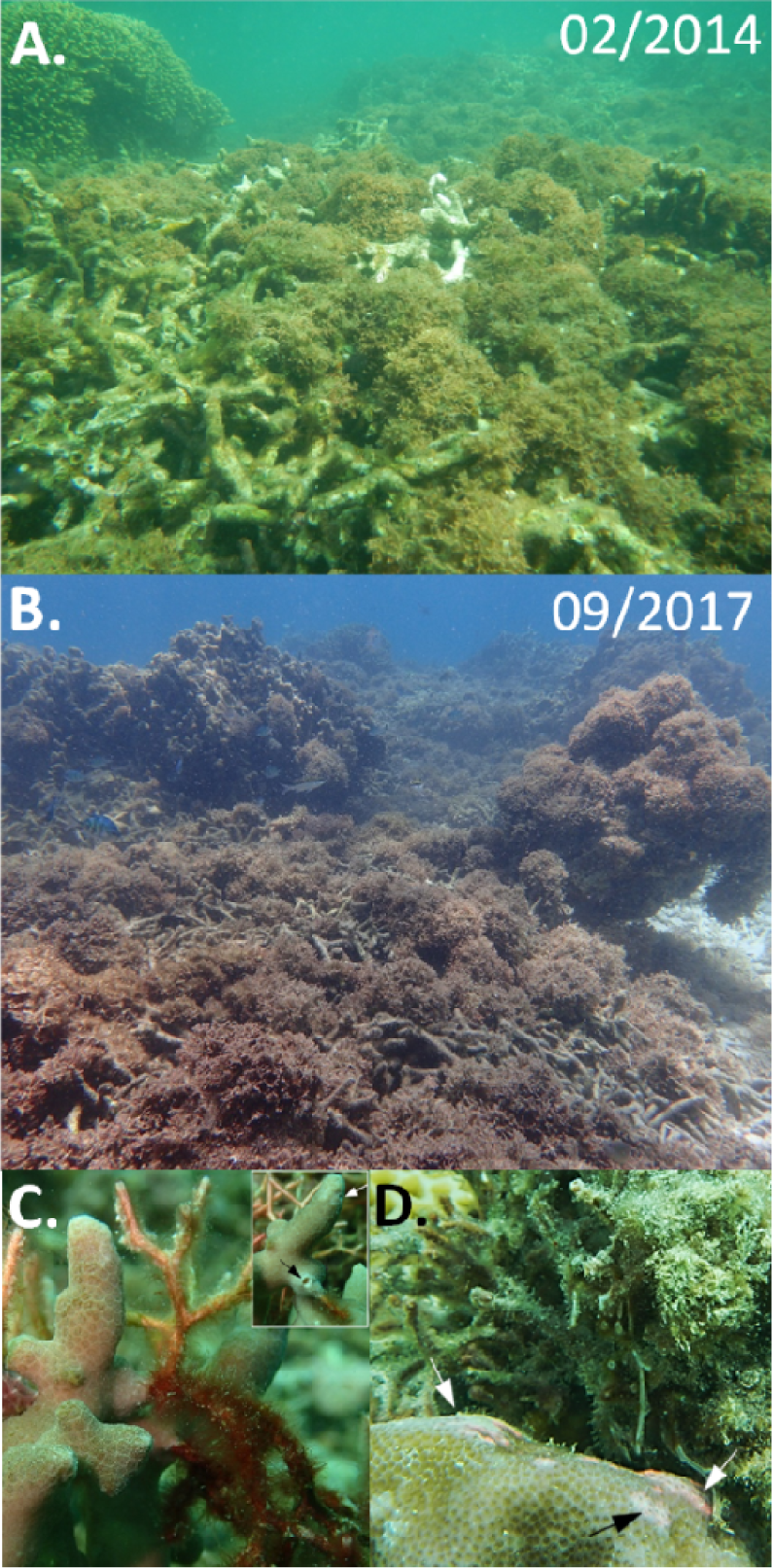
Observational photo-evidence of a prolonged *Galaxaura* overgrowth. A-B) A degraded patch reef in the southeast lagoon of Dongsha Atoll has been overgrown by *G. divaricata* for at last 3.5 years (3-5 m depth at site 9). Photos were taken in A) February 2014, with water temperature = 22.5°C; and B) in September 2017 with water temperature = 29°C. The holdfast of *G. divaricata* penetrates a branching *Porites* coral (*P. cylindrica*), creating small holes (inset). D) Coral (*P. solida*) tissue discoloration and bleaching (arrows) following direct contact with *G. divaricata*, potentially caused by allelopathic chemicals.

### Epiphytic macroalgae associated with *G. divaricata*

We identified 21 taxonomic groups of macroalgae, including macroscopic filamentous cyanobacteria, in association with *G. divaricata* (Table 1 and S2 Table). Among these, 15 were identified to the species level, with seven species of red algae, three species of brown, and five species of green algae (Table 1 and S2 Table). The most common green macroalgae associated with *G. divaricata* were *Derbesia marina* (occurrence frequency: 37%) (Fig. 6A), *Caulerpa chemnitzia* (27%) (Fig 6B), and *Boodlea composita* (20%). The most common brown macroalgae associated with *G. divaricata* were the brown algae *Lobophora* sp. (as *Lobphora* sp28 in [32]) (57%), *Padina* sp. (as *Padina* sp5 in [33]) (53%), and *Dictyota bartayresiana* (30%) (Fig 6C). The most common red macroalgae associated with *G. divaricata* were *Hypnea caespitosa* (100%) (Fig 6D), *Coelothrix irregularis* (87%), *Ceramium dawsoniia* (43%). Lastly, epiphytic macroscopic cyanobacteria (> 1cm in height) had an occurrence frequency of 17%. Among these epiphytic macroalgae we observed that the allelopathic *Lobophora* (identified as *Lobphora* sp28; S2 Table) was also found to frequently overgrow corals in the Dongsha lagoon (Fig 7 and S2 Fig).

**Table 1.**
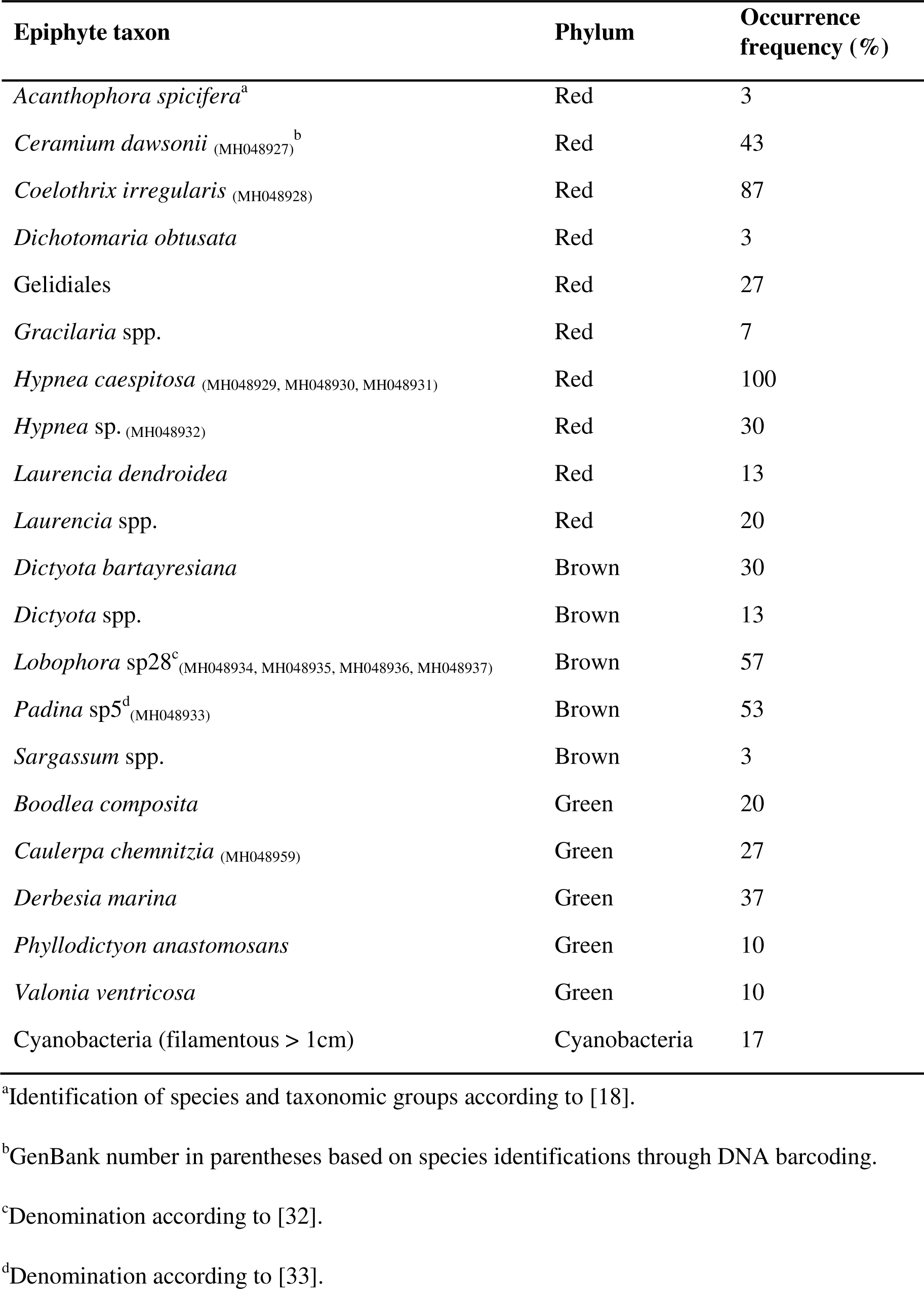
**Occurrence frequency (%) of epiphytic macroalgae on *Galaxaura divaricata* from the slope area of site 7.**

**Fig 6.**
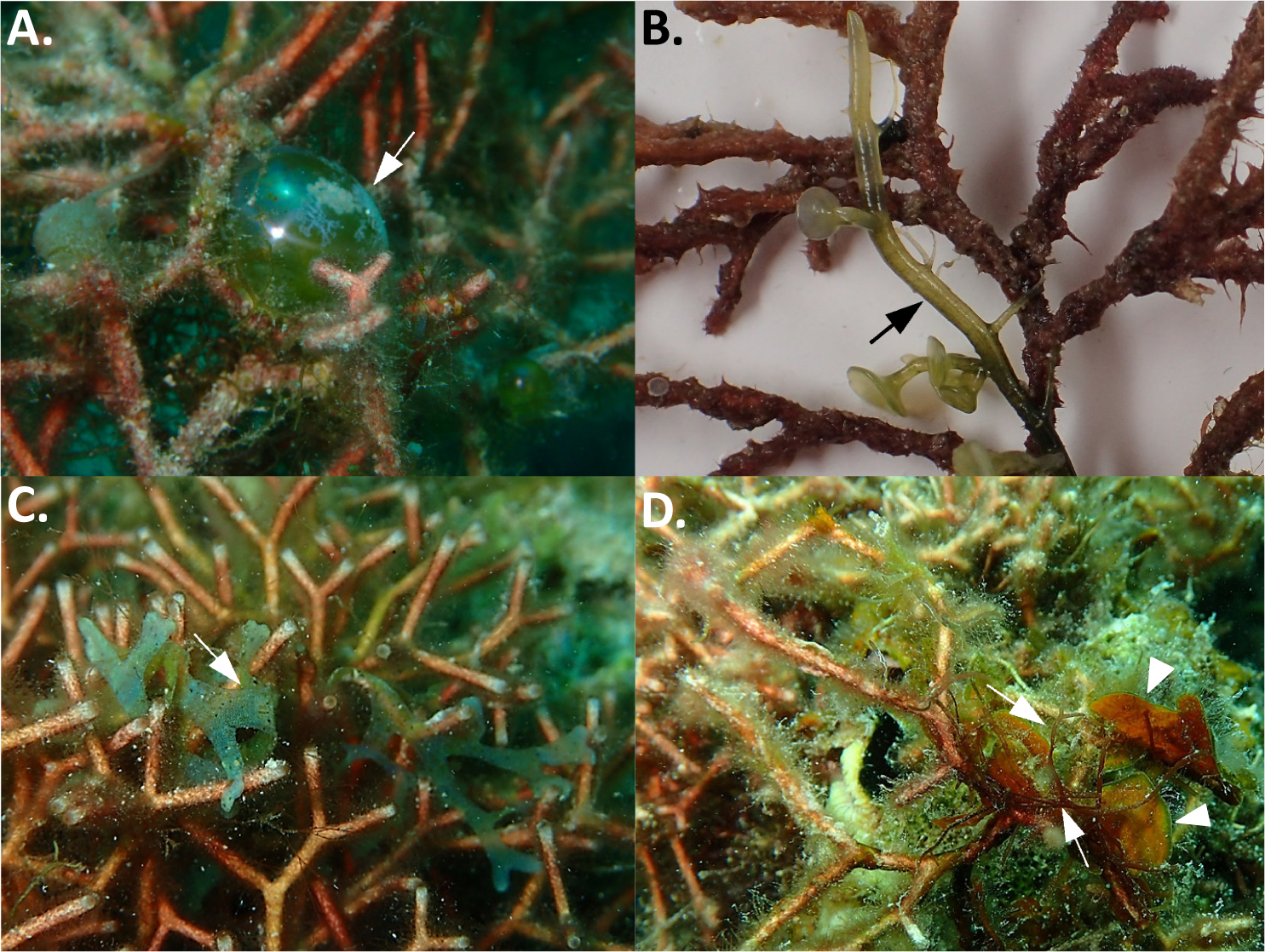
Examples showing epiphytic macroalgae that frequently grow on *G. divaricata*. A) *Valonia ventricosa*, B) *Caulerpa chemnitzia*, C) *Dictyota* sp., D) *Lobophora* sp. (arrowhead), and *Hypnea caespitosa* (arrow).

**Fig 7.**
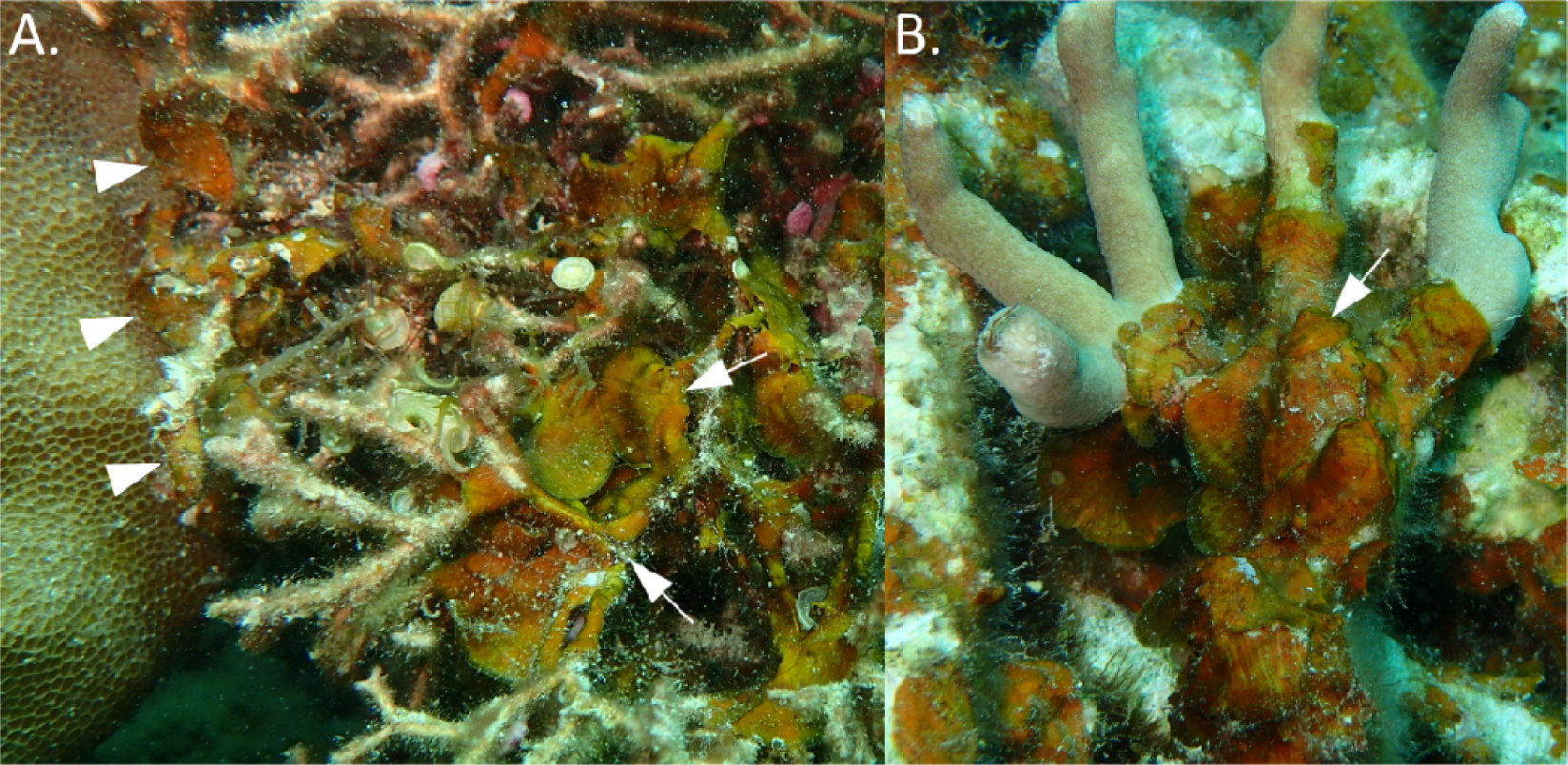
Coral overgrowth by *Lobophora* sp28. A) Example showing *Lobophora* sp28 growing on *Galaxaura divaricata* (arrows), and in contact with coral (*Porites solida*) (arrowheads). B) Coral overgrowth (*Porites cylindrica* in this case) by *Lobophora* sp28 is widespread in the shallow lagoon of Dongsha Atoll.

## Discussion

Our study shows that most patch reefs in the lagoon of Dongsha Atoll are degraded. Many reefs have a low live coral cover (below 25%) and high proportions of macroalgae, dead corals, and rubble, all of which are signs of reef degradation [34]. This is consistent with previous surveys that reported degraded conditions of lagoon patch reefs at Dongsha [35,36]. The filamentous form of *Galaxaura divaricata* overgrows degraded patch reefs in the southeast lagoon. This area is sheltered by a 2 km-wide reef flat, harboring shallow (1-5 m) and calm waters that may provide suitable growth conditions for *G. divaricata*. The proliferation of macroalgae is likely the consequence of an initial coral decline [37,38]. The synergistic effects of thermal stress, overfishing, and typhoon damage may have caused the decline of the once pristine corals in the Dongsha lagoon, followed by a proliferation of *G. divaricata*, among other macroalgae. Thermal stress on corals has increased over the past decades, with waters surrounding Dongsha Atoll warming at a faster rate than other areas of the South China Sea [35,39,40]. Recurrent bleaching events have caused high coral mortality and eradicated thermo-sensitive coral genera from the lagoon [41]. Overfishing and the extensive use of dynamite and cyanide, prior to the establishment of the Dongsha Atoll National Park in 2007 reduced fish, and destroyed large areas of coral framework [20,42]. Insufficient grazing after disturbance can lead to the establishment and full outgrowth of macroalgae beyond their initial stages [43]. *Galaxaura* is known to be largely unpalatable for various herbivorous fishes due to its calcareous thallus and low nutritional content [44–46]. Local herbivorous fish population in the Dongsha lagoon may not be effective to control the outgrowth of *Galaxaura* in certain areas.

Semi-closed lagoons are highly vulnerable to eutrophication and hypoxia, especially under the backdrop of climate change [47,48]. Reoccurring events of hypoxia during hot summers in 2014 and 2015 have caused substantial mass-die offs of the coral associated fauna and flora in the Dongsha lagoon [49]. Particularly, densities of macroinvertebrates, including echinoids, sea cucumbers, lobsters, and giant clams are extremely low (Table S4). *Galaxaura* appears to be well adapted to hypoxic conditions. For instance, *G. filamentosa* was one of the few algae to proliferate after a mass-die off caused by hypoxia in an atoll lagoon in French Polynesia [50].

Although the filamentous *G. divaricata* is a common allelopathic seaweed in subtropical and tropical waters, it has never been reported as a nuisance in overgrowing coral reefs. Our observations are the first to report a prolonged *G. divaricat*a overgrowth in degraded coral reefs. For instance, the *G. divaricata* cover was equally high on a degraded reef after 17-months. We further provide photo-evidence from another patch reef showing that the same *G. divaricata* overgrowth was present to a similar extend after 3.5 years. The photos clearly show that *G. divaricata* dominated the reef substrate of the site in both, the cooler northeast monsoon (winter) season (Fig 5.A, water temperature: 22.5°C), and the warmer southwest monsoon (summer) season (Fig 5.B, water temperature: 29°C). Due to challenging weather conditions, we were only able to conduct our quantitative temporal survey in April, the last month of the winter season, and therefore we cannot rule out potential variations in *G. divaricata* cover over the full length of that season. Expanding temporal surveys in the future will be worth of doing to confirm the long-term persistence of *G. divaricata* overgrowth.

A prolonged overgrowth of filamentous *G. divaricata* may have profound implications for the recovery potential of degraded reefs at Dongsha Atoll. Owning to its allelopathic effects on corals long-standing canopies of *G. divaricata* are likely to hamper coral recruitment ultimately preventing coral recovery [15,51]. As a caveat of this study, it is important to note that we did not attempt to isolate and identify allelopathic chemicals in *G. divaricata*. But, previous studies have identified lipid-soluble terpenoid compounds from filamentous *Galaxaura* cell extracts as allelochemicals that were capable of bleaching and killing of coral tissue [13]. It is also known that *Galaxaura* can change the chemical microclimate on degraded reefs with adverse effects on fish feeding behavior [4]. For instance, butterflyfish and other corallivores avoid corals in close association with *Galaxaura*, making it potentially difficult for these trophic guilds to find food [52,53]. Unlike other calcifying algae, such as coralline algae, *Galaxaura* does not stabilize the reef matrix. Thus, a prolonged *Galaxaura* overgrowth may contribute to the erosion and flattening of the reef structure, which negatively impacts biodiversity, and trophic support for coral associated organisms [54].

The filamentous *G. divaricata* is used as habitat by a variety of macroalgae. The availability of new habitat for epiphytic macroalgae provided by a prolonged *Galaxaura* overgrowth could have several implications for the ecology and recover potential of the reef. For instance, nutrient rich epiphytes could provide trophic support for herbivorous fishes and invertebrate, such as crustaceans and mollusks [24,55,56]. On the other hand, the association with the unpalatable *Galaxaura* may provide a refuge from herbivory for certain palatable algae [38,57], and facilitate their establishment on the reef, increasing macroalgae biodiversity [58]. The facilitation of harmful, allelopathic algal types could decrease the resilience and promote alternative stable states on coral reefs [59]. Some of the identified *G. divaricata* epiphytes, such as cyanobacteria [10], *Dictyota* [60], and *Lobophora* [9,61] are widely shown to overgrow corals after disturbance, and are known for their allelopathic inhibition of coral larvae recruitment. Here, we firstly report that an undescribed species *Lobophora* sp. (as *Lobophora* sp28 in [32]), the third most abundant macroalga on *G. divaricata*, overgrows and kills corals in the Dongsha lagoon through epizoism (Fig 7 and S2). Moreover, the microscopic filaments of *G. divaricata* may facilitate the attachment of macroalgae spores, while the calcified branches may provide structural support for fine, filamentous macroalgae. Considering that an increased substrate availability can promote macroalgae biomass on coral reefs, we hypothesize that, by providing a habitat for epiphytic macroalgae, *G. divaricata* may facilitate the diversity and abundance of macroalgae on degraded reefs. This study is merely observational and does not provide experimental evidence for the facilitation of macroalgae diversity and abundance by *G. divaricata.* However, the abovementioned hypotheses would be of great interest awaiting future validation.

## Conclusions

Our observations illustrated that the allelopathic and unpalatable filamentous seaweed, *Galaxaura divaricata*, can become dominant on degraded reefs in shallow, sheltered, and calm environments. We show that *G. divaricata* provides suitable substrate for a variety of macroalgae, further facilitating macroalgae growth and abundance on degraded reefs. Thus, a prolonged proliferation of *Galaxaura* could potentially enhance negative feedback loops, thereby perpetuating reef degradation. Several common epiphytic macroalgae on *Galaxaura* are allelopathic and known to frequently overgrow corals. Macroalgal assemblages, such as the *Galaxaura-*epiphyte system, warrant further investigation to better understand their ecological implications on the resilience of coral reefs, especially of shallow atoll lagoons. There are 439 listed coral reef atolls on earth; among them are 335 with semi-enclosed lagoons [62]. Atoll lagoons are highly productive and serve as valuable nursery habitat for marine life; however, they are most vulnerable to the effects of climate change [48,63]. Results from our study can be informative for the management and conservation of lagoons and shallow, inshore coral reef ecosystems, especially in the South China Sea and the Pacific Ocean, where filamentous *Galaxaura* is very common.

## Acknowledgements

The authors would like to thank our colleagues of the joint project: “Patterns of Resilience in Dongsha Atoll Coral Reefs” for their collaboration and great support throughout this study. We thank Keryea Soong and staff of the Dongsha Atoll Research Station, the Dongsha Atoll National Park, the Coastal Guard Administration, and the Ministry of Marine Affairs for logistic support. We would like to thank George P. Lohmann and Cherng-Shyang Chang for assistance with benthic surveys, as well as Chieh-Hsuan Lee and Pin-Chen Chen for assistance with fieldwork and DNA barcoding.

## Author contributions

### Conceptualization

Carolin Nieder, Shao-Lun Liu.

### Formal analysis

Carolin Nieder, Shao-Lun Liu.

### Investigation

Carolin Nieder.

### Writing–original draft

Carolin Nieder, Chaolun Allen Chen, Shao-Lun Liu.

### Writing–review & editing

Carolin Nieder, Chaolun Allen Chen, Shao-Lun Liu.

---

**S1 Fig.**
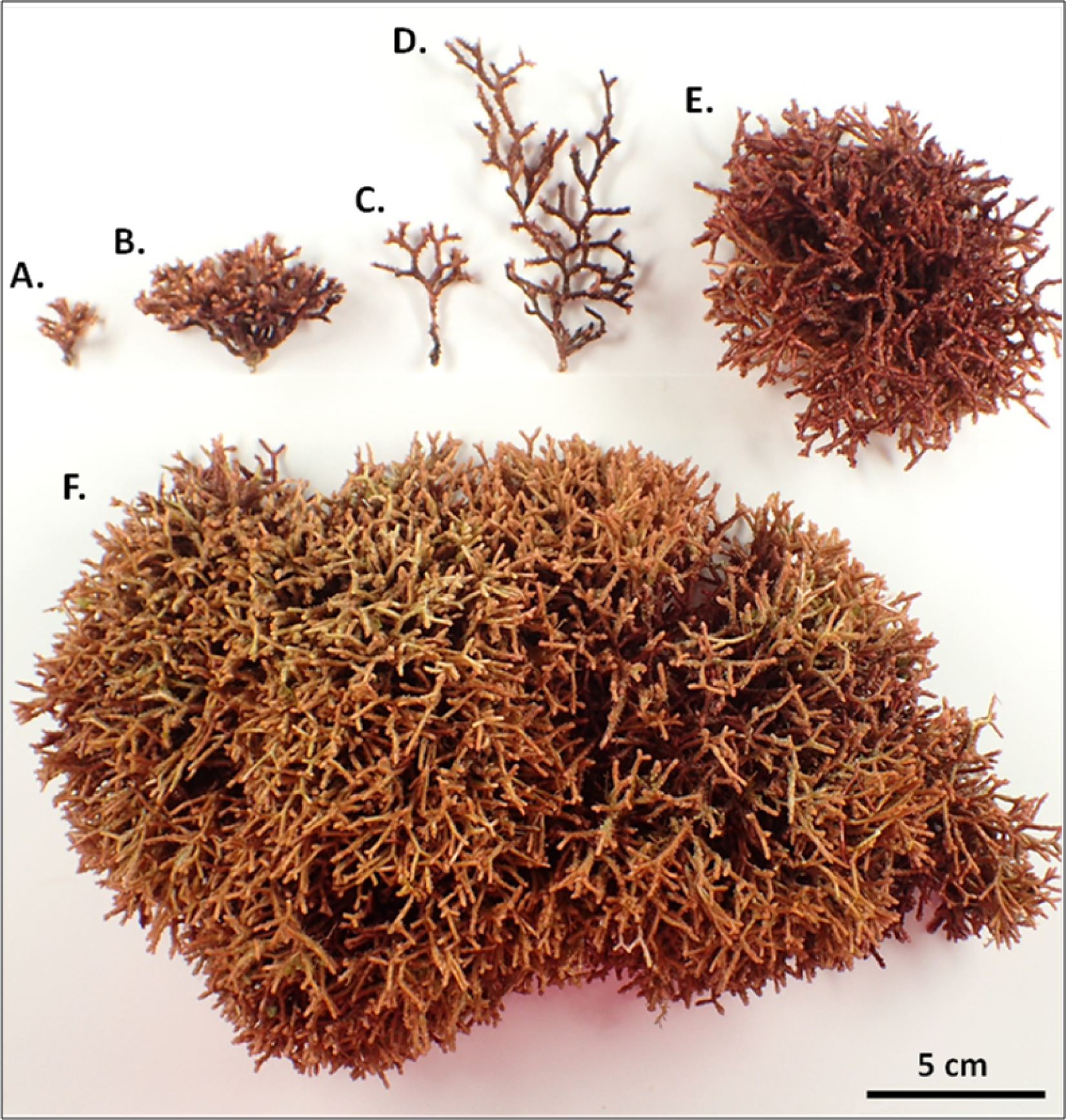
Various sizes and thallus shapes of *Galaxaura divaricata* from different locations in the lagoon of Dongsha Atoll. A-B) Small, ball-shaped thalli, and C-D) small, slender thalli were dominant on patch reefs in the north and northeast lagoon. E) Medium, ball-shaped thalli, and F) large carpet-like thalli were exclusively present in the southeast section of the lagoon.

**S2 Fig.**
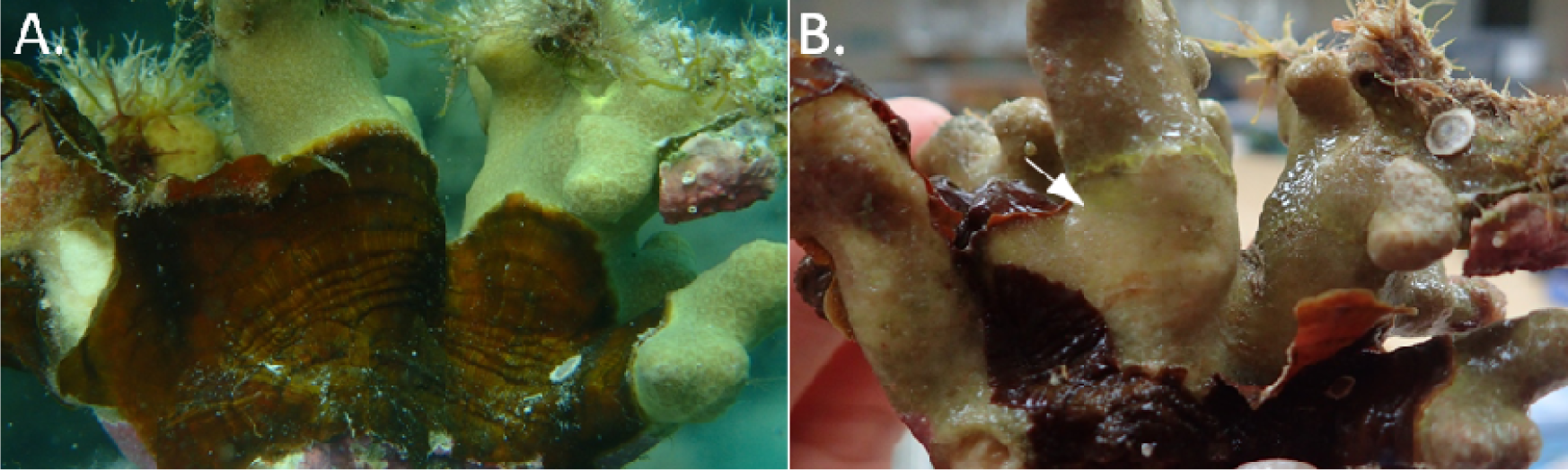
Coral overgrowth by *Lobophora* sp28. A) Coral overgrowth (*Porites cylindrica* in this case) by *Lobophora* sp28 is wide spread in the lagoon of Dongsha Atoll. B) The same coral showing dead tissue (arrow) after the removal of the algae.

**S1 Table.** **GPS coordinates of patch reef survey sites in the lagoon of Dongsha Atoll, South China Sea (Taiwan).**

| Site | Latitude | Longitude |
| --- | --- | --- |
| 1 | 20°44'26.28" | 116°47'8.879" |
| 2 | 20°43'31.62" | 116°47'52.679" |
| 3 | 20°42'16.14" | 116°48'27.419" |
| 4 | 20°44'20.52" | 116°51'35.699" |
| 5 | 20°42'29.88" | 116°52'54.599" |
| 6 | 20°42'6.72" | 116°50'53.279" |
| 7 | 20°38'24" | 116°50'20.999" |
| 8 | 20°38'3.12" | 116°49'30.479" |
| 9 | 20°36'52.86" | 116°49'24.179" |
| 10 | 20°36'53.4" | 116°46'2.399" |
| 11 | 20°39'52.2" | 116°46'32.159" |
| 12 | 20°41'49.2" | 116°44'45.659" |
| 13 | 20°42'12.36" | 116°44'14.639" |

**S2 Table.** **Information and Genbank numbers of macroalgae samples used for DNA barcoding in this study.**

| Species | Phylum | Voucher# | Date | Site | Area | Substrate | Marker | GenBank# |
| --- | --- | --- | --- | --- | --- | --- | --- | --- |
| *Ceramium dawsonii* | Red | KO208 | Oct-16 | 7 | slope | *G. divaricata* | *rbcL* | MH048927 |
| *Coelothrix irregularis* | Red | K0206 | Oct-16 | 7 | slope | *G. divaricata* | *rbcL* | MH048928 |
| *Hypnea caespitosa* | Red | K0204 | Oct-16 | 7 | slope | *G. divaricata* | *rbcL* | MH048929 |
| *Hypnea caespitosa* | Red | K0203 | Oct-16 | 7 | slope | *G. divaricata* | *rbcL* | MH048930 |
| *Hypnea caespitosa* | Red | SD17120 | Aug-17 | 7 | slope | *G. divaricata* | *rbcL* | MH048931 |
| *Hypnea* sp. | Red | K0205 | Oct-16 | 7 | slope | *G. divaricata* | *rbcL* | MH048932 |
| *Lobophora* sp28<sup>a</sup> | Brown | SD17023 | Aug-17 | 7 | slope | *G. divaricata* | *rbcL* | MH048934 |
| *Lobophora* sp28 | Brown | SD17021 | Aug-17 | 7 | slope | *G. divaricata* | *rbcL* | MH048935 |
| *Lobophora* sp28 | Brown | SD17019 | Aug-17 | 7 | slope | *G. divaricata* | *rbcL* | MH048936 |
| *Lobophora* sp28 | Brown | SD17017 | Aug-17 | 7 | slope | *G. divaricata* | *rbcL* | MH048937 |
| *Lobophora* sp28 | Brown | K0173 | Apr-16 | 7 | slope | Coral | *rbcL* | MH048940 |
| *Lobophora* sp28 | Brown | SD17058 | Aug-17 | 9 | slope | Coral | *rbcL* | MH048941 |
| *Padina* sp5<sup>b</sup> | Brown | SD17114 | Aug-17 | 7 | slope | *G. divaricata* | *rbcL* | MH048933 |
| *Caulerpa chemnitzia* | Green | SD17117 | Aug-17 | 7 | slope | *G. divaricata* | *tufA* | MH048959 |

<sup>a</sup>Denomination according to [1].
<sup>b</sup>Denomination according to [2].

**S3 Table.** **Information and Genbank numbers of *Galaxaura divaricata* samples from various locations in the lagoon of Dongsha Atoll that were used for DNA barcoding in this study.**

| Species | Voucher# | Date | Site | Area | Substrate | Thallus size | Marker | GenBank# |
| --- | --- | --- | --- | --- | --- | --- | --- | --- |
| *G. divaricata* | K0210 | Apr-16 | 7 | slope | rock | medium | *rbcL* | MH048946 |
| *G. divaricata* | R90B12 | Feb-14 | 9 | top | rock | large | *rbcL* | MH048942 |
| *G. divaricata* | SD17048 | Aug-17 | 9 | slope | rubble | large | *rbcL* | MH048957 |
| *G. divaricata* | SD17098 | Aug-17 | 6 | top | rock | small | *rbcL* | MH048943 |
| *G. divaricata* | SD17099 | Aug-17 | 1 | top | rock | small | *rbcL* | MH048956 |
| *G. divaricata* | SD17100 | Aug-17 | 5 | top | rock | small | *rbcL* | MH048955 |
| *G. divaricata* | SD17101 | Aug-17 | 5 | top | rock | small | *rbcL* | MH048958 |
| *G. divaricata* | SD17102 | Aug-17 | 5 | slope | rock | medium | *rbcL* | MH048954 |
| *G. divaricata* | SD17103 | Aug-17 | 1 | slope | rock | small | *rbcL* | MH048953 |
| *G. divaricata* | SD17104 | Aug-17 | 6 | slope | rock | medium | *rbcL* | MH048952 |
| *G. divaricata* | SD17105 | Aug-17 | 5 | slope | rock | medium | *rbcL* | MH048951 |
| *G. divaricata* | SD17106 | Aug-17 | 6 | slope | rock | small | *rbcL* | MH048950 |
| *G. divaricata* | SD17107 | Aug-17 | 6 | slope | rock | small | *rbcL* | MH048944 |
| *G. divaricata* | SD17110 | Aug-17 | 6 | top | coral | medium | *rbcL* | MH048949 |
| *G. divaricata* | SD17112 | Aug-17 | 4 | slope | rock | medium | *rbcL* | MH048948 |
| *G. divaricata* | SD17113 | Aug-17 | 4 | slope | rock | small | *rbcL* | MH048947 |

**S4 Table.** **Statistics summary of 2-way ANOVA of major benthic categories (live corals, total macroalgae, CCA, and other substrates) between two reef areas among 12 sites.**

| Source of variation | *df* | Mean Sq | *F* | *P* |
| --- | --- | --- | --- | --- |
| *Corals* |  |  |  |  |
| area (top > slope; dif: 3.15) | 1 | 2673 | 6.617 | 0.0102* |
| site | 11 | 11036 | 27.321 | <2e-16* |
| area × site | 11 | 7110 | 17.601 | <2e-16* |
| error | 1056 | 404 |  |  |
| *Total macroalgae* |  |  |  |  |
| area (top > slope; dif: 5.03) | 1 | 6835 | 20.68 | 6.07e-06* |
| site | 11 | 16025 | 48.48 | <2e-16* |
| area × site | 11 | 8684 | 26.27 | <2e-16* |
| error | 1056 | 331 |  |  |
| *Crustose coralline algae (CCA)* |  |  |  |  |
| area (top > slope; dif: 1.36) | 1 | 500.2 | 62.725 | 5.99e-15* |
| site | 11 | 44.0 | 5.514 | 1.29e-08* |
| area × Site | 11 | 28.8 | 3.616 | 4.88e-05* |
| error | 1056 | 8.0 |  |  |
| *Other substrates* |  |  |  |  |
| area (top < slope; dif: 9.54) | 1 | 24548 | 41.30 | 1.97e-10* |
| site | 11 | 18205 | 30.63 | <2e-16* |
| area × site | 11 | 10405 | 17.51 | <2e-16* |
| error | 1056 | 594 |  |  |

\*Asterisk indicates a statistically significance at the p value < 0.05.

**S5 Table.** **Statistics summary of 2-way ANOVA of the *G. divaricata* percent cover between two reef areas among 13 sites.**

| Source of variation | *df* | Mean Sq | *F* | *P* |
| --- | --- | --- | --- | --- |
| *G. divaricata* |  |  |  |  |
| area (top < slope; dif: 1.20) | 1 | 423 | 6.574 | 0.0105* |
| site | 12 | 11536 | 179.278 | <2e-16* |
| area × site | 12 | 480 | 7.460 | 2.28e-13* |
| error | 1144 | 64 |  |  |

\*Asterisk indicates a statistically significance at the p value < 0.05.

**S6 Table.** **Percent cover of *Galaxaura divaricata* on 13 patch reef sites in the lagoon of Dongsha Atoll, South China Sea.**

| Site | Location | Patch reef area |  | Cover rank |
| --- | --- | --- | --- | --- |
|  |  | Top (1-5 m) | Slope (5-10 m) |  |
| 1 | North | 0.02 ± 0.08<sup>a</sup> | 0.02 ± 0.15 | very low |
| 2 | North | 0 | 0 | very low |
| 3 | North | 0.27 ± 0.78 | 0.31 ± 0.9 | very low |
| 4 | Northeast | 1.52 ± 1.9 | 1.47 ± 1.95 | very low - low |
| 5 | Northeast | 5.69 ± 5.47 | 3.37 ± 5.62 | low - moderate |
| 6 | Northeast | 0.79 ± 1.36 | 0.21 ± 0.34 | very low |
| 7 | Southeast | 0.02 ± 0.15 | 15.86 ± 17 | very low - high |
| 8 | Southeast | 0.17 ± 0.67 | 4.31 ± 6.67 | very low - low |
| 9 | Southeast | 41.87 ± 24.73 | 40.87 ± 25.58 | high |
| 10 | South | 0 | 0 | very low |
| 11 | Center | 0.46 ± 1.41 | 0.02 ± 0.15 | very low |
| 12 | West | 0 | 0 | very low |
| 13 | West | 0 | 0 | very low |

<sup>a</sup>Data are relative cover expressed as mean ± SD (%) of 45 replicate quadrats for reef top and reef slope.

**S7 Table.** **Statistics summary of 2-way ANOVA of the percent cover in two major benthic categories (*G. divaricata* and corals) over time.**

| **Source of variation** | ***df*** | **Mean Sq** | ***F*** | ***P*** |
| --- | --- | --- | --- | --- |
| *Two benthic categories* |  |  |  |  |
| benthic category | 1 | 26.84 | 0.086 | 0.770 |
| time | 3 | 197.58 | 0.632 | 0.595 |
| benthic category × time | 3 | 113.44 | 0.363 | 0.780 |
| error | 352 | 312.61 |  |  |

**S8 Table.** **Paucity of macrobenthic invertebrates in the Dongsha lagoon.**

| Macrobenthic fauna | Count | Density (individual/100 m<sup>2</sup>) |
| --- | --- | --- |
| *Diadema savignyi* | 3 | 0.03 |
| *Diadema setosum* | 1 | 0.01 |
| *Echinometra mathaei* | 126 | 1.26 |
| *Echinothrix calamaris* | 3 | 0.03 |
| *Tripneustes gratilla* | 4 | 0.04 |
| *Culcita novaeguineae* | 54 | 0.54 |
| *Echinaster luzonicus* | 2 | 0.02 |
| *Fromia* spp. | 8 | 0.08 |
| *Linckia multifora* | 15 | 0.15 |
| *Holothuria* | 0 | 0 |
| *Cypraea tigris* | 3 | 0.03 |
| Giant clam | 34 | 0.34 |
| *Lambis* spp. | 3 | 0.03 |
| Lobster | 0 | 0 |

Data derived from a belt transect survey of 13 patch reefs and seven seagrass beds (10,000 m<sup>2</sup> total area surveyed) across the lagoon of Dongsha Atoll in September 2017.

